# Convergence of peptidergic and non-peptidergic protein markers in the human dorsal root ganglion and spinal dorsal horn

**DOI:** 10.1101/2020.10.14.339382

**Authors:** Stephanie Shiers, Ishwarya Sankaranarayanan, Vivek Jeevakumar, Anna Cervantes, Jeffrey C. Reese, Theodore J. Price

**Affiliations:** University of Texas at Dallas, School of Behavioral and Brain Sciences and Center for Advanced Pain Studies; Southwest Transplant Alliance

**Keywords:** DRG, dorsal horn, CGRP, TRPV1, Nav1.7, P2X3

## Abstract

Peripheral sensory neurons are characterized by their size, molecular profiles, and physiological responses to specific stimuli. In mouse, the peptidergic and non-peptidergic subsets of nociceptors are distinct and innervate different lamina of the spinal dorsal horn. The unique molecular signature and neuroanatomical organization of these neurons supports a labeled line theory for certain types of nociceptive stimuli. However, long standing evidence supports the polymodal nature of nociceptors in many species. We have recently shown that the peptidergic marker, CGRP, and the non-peptidergic marker, P2X3R, show largely overlapping expression at the mRNA level in human dorsal root ganglion (DRG). Herein, our aim was to assess the protein distribution of nociceptor markers, including their central projections, in the human DRG and spinal cord. Using DRGs obtained from organ donors, we observed that CGRP and P2X3R were co-expressed by approximately 33% of human DRG neurons and TrpV1 was expressed in ~60% of human DRG neurons. In the dorsal spinal cord, CGRP, P2X3R, TrpV1 and Nav1.7 protein stained the entirety of lamina II, with only P2XR3 showing a gradient of expression. This was confirmed by measuring the size of the substantia gelatinosa using Hematoxylin and Eosin staining of adjacent sections. Our findings are consistent with the known polymodal nature of most primate nociceptors and indicate that the central projection patterns of nociceptors are different between mice and humans. Elucidating how human nociceptors connect to subsets of dorsal horn neurons will be important for understanding the physiological consequences of these species differences.

## Introduction

Nociceptors found in the dorsal root ganglia (DRG) are responsible for transmitting nociceptive information to the dorsal horn of the spinal cord. In mouse, these neurons have been classified into multiple subtypes based on their size, neurochemical signatures, gene expression profiles, and their functional responsiveness to thermal, mechanical and chemical stimuli. These subpopulations form synapses on central neurons of the dorsal horn in an organized manner throughout lamina I, II and V (Abrahamsen et al., 2008; Ribeiro-da-Silva, 2015). Calcitonin gene-related peptide (CGRP) and P2X purinergic ion channel type 3 receptor (P2X3R) have been identified at the mRNA and protein levels to be expressed in distinct populations of mouse DRG neurons using an array of new and old technologies including single-cell sequencing (Abrahamsen et al., 2008; Shiers, Klein, & Price, 2020; Usoskin et al., 2015). These markers have been extensively utilized to distinguish peptidergic (CGRP/substance P-releasing) and non-peptidergic (P2X3R-expressing or isolectin B4-binding (IB4)) nociceptors in the mouse DRG and are also used in the rat, although they show overlap in this species (T. J. Price & Flores, 2007). These peptidergic and non-peptidergic neurons form distinct synaptic connections in the mouse spinal dorsal horn with CGRP afferents being found primarily in lamina I and lamina IIo while P2X3R/IB4 positive afferents are almost exclusively localized to lamina IIi (Abrahamsen et al., 2008; Eftekhari & Edvinsson, 2011; Fan, Kim, Warner, & Gustafsson, 2007; Park et al., 2008). These nociceptor projections have not been quantified in the human dorsal horn, but previous histochemical studies in human and non-human primates suggest that peptidergic nociceptors project uniformly throughout lamina II in these species (Carlton, McNeill, Chung, & Coggeshall, 1987; Harmann, Chung, Briner, Westlund, & Carlton, 1988; McNeill, Carlton, & Hulsebosch, 1991; Pawlowski et al., 2013).

The existence in mouse of distinct nociceptor subtypes and their structured organization in the spinal dorsal horn supports the idea of modality-based stimulus encoding, a “labeled line” for certain types of behavioral nociceptive and pruritic responses (J. Braz, Solorzano, Wang, & Basbaum, 2014; J. M. Braz, Nassar, Wood, & Basbaum, 2005; Cavanaugh et al., 2009; Huang et al., 2019; Mishra & Hoon, 2013). However, the polymodal nature of most nociceptors across species has long been recognized (Perl, 1996). A potential explanation of these disparate findings are species differences in nociceptor subsets and their central projections. In support of this, we recently showed that there is very little distinction between CGRP and P2X3R mRNA-expressing neurons in the human DRG (Shiers et al., 2020). Previous work shows that these nociceptor populations also overlap in rats (T. J. Price & Flores, 2007), albeit to a lesser extent. This blurred peptidergic/non-peptidergic signature in the human DRG is supported by other striking species differences (Shiers et al., 2020). For example, in mouse, the transient receptor potential cation channel subfamily V member 1 (TrpV1) has been identified at the protein and mRNA level to be expressed in ~30% of peripheral sensory neurons. Ablation of TrpV1 neurons in mice specifically occludes thermal pain sensitivity suggesting that this neuronal population encodes a labeled line for noxious heat (Abrahamsen et al., 2008). In mouse spinal cord, TrpV1 afferents are primarily localized to lamina I and lamina IIo of the dorsal horn, similar to CGRP (Cavanaugh et al., 2009). In rat there is evidence that these projections are different as much TrpV1 immunoreactivity in the rat spinal cord overlaps with IB4 staining (Guo et al., 2001; Guo, Vulchanova, Wang, Li, & Elde, 1999). Unlike rodents, most human nociceptors produce mRNA for *TRPV1* (Shiers et al., 2020), a finding that is consistent with microneurography recordings wherein most human nociceptors respond to capsaicin (Schmelz, Schmid, Handwerker, & Torebjork, 2000). Our goal in the work described here was to thoroughly characterize protein expression for nociceptor markers in the human DRG and dorsal horn with the purpose of providing information needed to understand species differences in nociceptor populations and their central projections.

We used immunohistochemistry (IHC) for CGRP, P2X3R, TrpV1 and Nav1.7 to describe their distribution in the human DRG and dorsal spinal cord. We find that their protein profiles are very similar to previously reported mRNA expression patterns in human DRG (Shiers et al., 2020). In the dorsal horn these proteins show diffuse neuropil staining throughout the entire substantia gelatinosa, encompassing all of lamina II. Only P2XR3 showed a gradient of expression. Our work supports the conclusion that there are fundamental differences in nociceptor populations between mice and humans and that these differences are represented in the central projections of these neurons.

## Materials and Methods

### Tissue preparation

All human tissue procurement procedures were approved by the Institutional Review Boards at the University of Texas at Dallas. Human lumbar dorsal root ganglion and spinal cord were collected, frozen on dry ice and stored in a −80°C freezer. Donor information is provided in **Table 1**. Tissues were collected from organ donors within 2 hours of cross-clamp. The human DRGs and spinal cords were gradually embedded in OCT in a cryomold by adding small volumes of OCT over dry ice to avoid thawing. All tissues were cryostat sectioned at 20 μm onto SuperFrost Plus charged slides. Sections were only briefly thawed in order to adhere to the slide but were immediately returned to the −20°C cryostat chamber until completion of sectioning. The slides were then immediately utilized for histology.

**Table 1.** Human donor information. Donor information is given for all the samples that were used in all experiments. STA = Southwest Transplant Alliance

| Donor | DRG | Spinal cord | Sex | Age | Cause of death | Collection site |
| --- | --- | --- | --- | --- | --- | --- |
| 1 | Lumbar | Lumbar | Male | 29 | Head trauma | STA |
| 2 | Lumbar | Lumbar | Female | 44 | Cardiac arrest | STA |
| 3 | Lumbar | Lumbar | Female | 62 | Stroke | STA |
| 4 | Lumbar | Lumbar | Male | 43 | Head Trauma | STA |
| 5 | Lumbar |  | Female | 33 | Opioid overdose | STA |
| 6 | Lumbar 5 |  | Female | 48 | Head Trauma | Anabios |

### Immunohistochemistry (IHC)

Slides were removed from the cryostat and immediately transferred to cold 10% formalin (4°C; pH 7.4) for 15 minutes. The tissues were then dehydrated in 50% ethanol (5 min), 70% ethanol (5 min), 100% ethanol (5 min), 100% ethanol (5 min) at room temperature. The slides were air dried briefly and then boundaries were drawn around each section using a hydrophobic pen (ImmEdge PAP pen, Vector Labs). When hydrophobic boundaries had dried, the slides were submerged in blocking buffer (10% Normal Goat Serum, 0.3% Triton-X 100 in 0.1M Phosphate Buffer (PB)) for 1 hour at room temperature. Slides were then rinsed in 0.1M PB, placed in a light-protected humidity-controlled tray and incubated in primary antibody diluted in blocking buffer overnight at 4°C. A list of all primary and secondary antibodies is shown in **Table 2**. The next day, slides were washed in 0.1M PB and then incubated in their respective secondary antibody (1:2000) with DAPI (1:5000; Cayman Chemical; Cat # 14285) diluted in blocking buffer for 1 hour at room temperature. The sections were washed in 0.1M PB and then covered with Sudan Black B (0.05% diluted in 70% Ethanol), a blocker of lipofuscin, for 10 minutes. The Sudan Black B step was performed in all IHC experiments except the CGRP/P2X3R human DRG experiment (due to issues with diminished quality in IHC label). Sections were then washed in 0.1M PB, air dried and coverslipped with Prolong Gold Antifade reagent.

**Table 2.** List of antibodies used for immunohistochemistry.

| Antibody | Vendor | Catalog # | RRID | Dilution/Conc. |
| --- | --- | --- | --- | --- |
| Rabbit-anti-CGRP | ImmunoStar | 24112 | AB_572217 | 1:1000 |
| Rabbit-anti-TrpV1 | ThermoFisher | PA1-748 | AB_2209010 | 1:500 |
| Mouse-anti-Nav1.7 | NeuroMab | N68/6 | AB_2184355 | 2µg/mL |
| Mouse-anti-P2X3 | Santa Cruz | B-5 | AB_2810268 | 5µg/mL |
| Chicken-anti-peripherin | Encor | CPCA-Peri | AB_228443 | 1:1000 |
| Goat-anti-rabbit H&L 555 | ThermoFisher | A21428 | AB_2535849 | 1:2000 |
| Goat-anti-rabbit H&L 488 | ThermoFisher | A11034 | AB_2576217 | 1:2000 |
| Goat-anti-chicken H&L 488 | ThermoFisher | A11039 | AB_2534096 | 1:2000 |
| Goat-anti-chicken H&L 647 | ThermoFisher | A21449 | AB_2535866 | 1:2000 |
| Goat-anti-mouse IgG1 488 | ThermoFisher | A21121 | AB_2535764 | 1:2000 |
| Goat-anti-mouse IgG1 555 | ThermoFisher | A21127 | AB_2535769 | 1:2000 |

### Antibody Characterization

The CGRP antibody (ImmunoStar, Cat # 24112, RRID: AB_572217) is a polyclonal antibody raised in rabbit and made against synthetic rat CGRP (SCNTATCVTHRLAG LLSRSGGVVKDNFVPTNVGSEAF-NH2). According to the manufacturer, CGRP immunolabeling was completely abolished by soluble pre-adsorption with rat CGRP at a final concentration of 10-5M. It has been extensively utilized for immunohistochemistry (Pubmed search: 75 papers) on mouse, rat, and human nervous tissue, particularly in human nerve (Donadio et al., 2019; Karlsson et al., 2020).

The P2X3R antibody (Santa Cruz, Cat # B-5, RRID: AB_2810268) is a mouse monoclonal antibody against amino acids 338-397 of the human P2X3R C-terminus. According to the manufacturer’s datasheet, the antibody recognizes the expected monomeric (44 kDa) band for P2X3R on a western blot of various whole cell lysates. It also has been published to show western blot specificity in human muscle afferents (Smith et al., 2020).

The TrpV1 antibody (ThermoFisher, Cat # PA1-748), RRID: AB_2209010) is a polyclonal antibody raised in rabbit against a synthetic human TrpV1 peptide (T(7) D L G A A A D P L Q K D T C(21). The antibody has been extensively utilized in western blot and IHC experiments on human tissues (de Fontgalland, Brookes, Gibbins, Sia, & Wattchow, 2014; Dinis et al., 2005; Gonzales et al., 2014; Pecze et al., 2008; Xiao, Zhou, Liu, Xie, & Guo, 2019). The antibody only recognizes the expected TrpV1 band (94 kDa) on a western blot of human trigeminal ganglion (Pecze et al., 2008).

The Nav1.7 antibody (NeuroMab, Cat # N68/6, RRID: AB_2184355) is a mouse monoclonal raised against amino acids 1751-1946 of the human Nav1.7 C-terminus. According to the manufacturer’s datasheet, the antibody recognizes the expected monomeric (230 kDa) band for Nav1.7 on cell lysates of transfected HEK cells and does not cross-react with other Nav channels. The antibody has been knockout-validated using immunocytochemistry on mouse cultured DRG neurons and using IHC on rat brain (Grubinska et al., 2019). We have also recently shown that this antibody robustly stains human DRG, and shows a specific and similar expression pattern to its mRNA (Shiers et al., 2020).

The peripherin antibody (EnCor, Cat # CPCA-Peri, RRID: AB_228443) is a chicken polyclonal raised against full-length recombinant rat peripherin. The manufacturer datasheet shows an intensely immunolabeled WB band at the desired molecular weight (57 kDa) on lysates of rat, mouse, and pig spinal cord.

### Hematoxylin and Eosin stain (H&E)

H&E staining was performed on adjacent human lumbar spinal cord sections to better visualize the substantia gelatinosa and other anatomy to compare to our IHC staining. After cryosectioning, slides were removed from the cryostat and immediately transferred to cold 10% formalin (4°C; pH 7.4) for 15 minutes. The tissue was then dehydrated in 50% ethanol (5 min), 70% ethanol (5 min), 100% ethanol (5 min), 100% ethanol (5 min) at room temperature. Isopropanol was pipetted onto each section and incubated for 1 minute at room temperature. The slides were then air dried briefly. Hematoxylin (Agilent Technologies; Cat S330930-2) was pipetted onto each section until completely covered and incubated for 7 minutes. The reagent was discarded and then the slide was washed three times in Milli-Q water. Bluing buffer (Agilent Technologies; Cat # CS703230-2) was then added to the slide until the sections were completely covered and incubated for 2 minutes at room temperature. The slides were washed in Milli-Q water and then Eosin mix (Eosin Y solution; Millipore Sigma; Cat #HT110216 with Tris-Acetic Acid Buffer) was added to each section and incubated for 1 minute at room temperature. The slides were then washed again in Milli-Q water and then allowed to air dry before being coverslipped with glycerol.

### Imaging

For all immunohistochemistry experiments, sections were imaged on an Olympus FV3000 confocal microscope at 10X magnification. For human DRGs, 2 10X images were acquired of each section, and 2-3 sections were imaged per human donor. For human spinal cord, 1 10X image of the dorsal horn was acquired per human donor. The acquisition parameters were set based on guidelines for the FV3000 provided by Olympus. In particular, the gain was kept at the default setting 1, HV ≤ 600, offset = 4, and laser power ≤ 10% (but generally the laser power was ≤ 5% for our experiments). All images were acquired using the same image settings. There was a single IHC experiment of human spinal cord (Nav1.7/Peripherin as shown in **Figure X**) that was imaged at 20x on an Olympus vs120 slide scanner in order to acquire a mosaic image of the entire section.

For H&E stained sections, brightfield images were acquired using an Olympus vs120 slide scanner at 20X magnification.

### Images Analysis

For human DRG IHC quantification, the raw images were brightened and contrasted equally in Olympus CellSens (v1.18), and then the fluorescence intensity and the diameter of each neuron was measured manually one neuron at a time for each target. The corrected fluorescence intensity value was calculated by subtracting the average fluorescence intensity signal acquired from all neurons in the negative control (exposed only to secondary antibody). All antibodies gave a gradient of signal in most neurons. Therefore, to calculate the strongest positives, we first determined the range of signal (fluorescence intensity) of all neurons for each section, and then divided it by 5 to define very low, low, moderate, high, and very high expressors. We considered all neurons positive for CGRP, TRPV1, or P2X3R if they gave a fluorescence intensity value in the moderate-high-very high expression range. Total neuron counts were acquired by counting all of the antibody-labeled neurons and all neurons that were clearly outlined by DAPI (satellite cell).

### Data Analysis

Graphs were generated using GraphPad Prism version 8.01 (GraphPad Software, Inc. San Diego, CA USA). A relative frequency distribution histogram with a fitted Gaussian distribution curve was generated using the diameters of target-positive neurons.

## Results

### CGRP and P2X3R protein show substantial overlap in human DRG

We first conducted immunohistochemistry on human DRG to gauge protein expression in human DRG neuronal populations. We used a human-specific CGRP antibody that has been characterized previously on human nervous system tissue (Donadio et al., 2019; Karlsson et al., 2020) and a human-specific P2X3R monoclonal antibody that shows a clean Western blot signal in characterization experiments (**Table 2**). It is important to note that neither of these antibodies have been knockout-validated; however, mouse-knockout validation does not necessarily confirm specificity in human. We observed a gradient of signal ranging from very low – very high signal for both antibodies (**Fig 1A**). We used a conservative quantitative approach and only classified neurons as positive for the protein target of the antibody when labelled neurons gave moderate to very high signal. Pie-chart representations of CGRP and P2X3R subpopulations show that there was very little variability between donors in the overall number of neurons labelled (**Fig 1B**). A small percentage of neurons expressed only CGRP (average 17.3%) or only P2X3R (13.8%) separately, but there was a larger overlapping population that expressed both proteins (average 33.1%) (**Fig 1B**). The total percentage of neurons expressing CGRP was 50.4% and for P2X3R was 48.3% (**Fig 1C**). In the images shown in **Fig 1**, most neurons appear yellow due to the colocalization of CGRP and P2X3R. The size profile of CGRP and P2X3R neurons indicated that these neurons are small-to-medium in size (**Fig 1D-E**). These findings are consistent with our previously published work on *CALCA* and *P2X3R* mRNA in human DRGs in which *CALCA* and *P2X3R* mRNA was present in ~60% and ~55% of all sensory neurons, respectively (Shiers et al., 2020).

**Figure 1.**
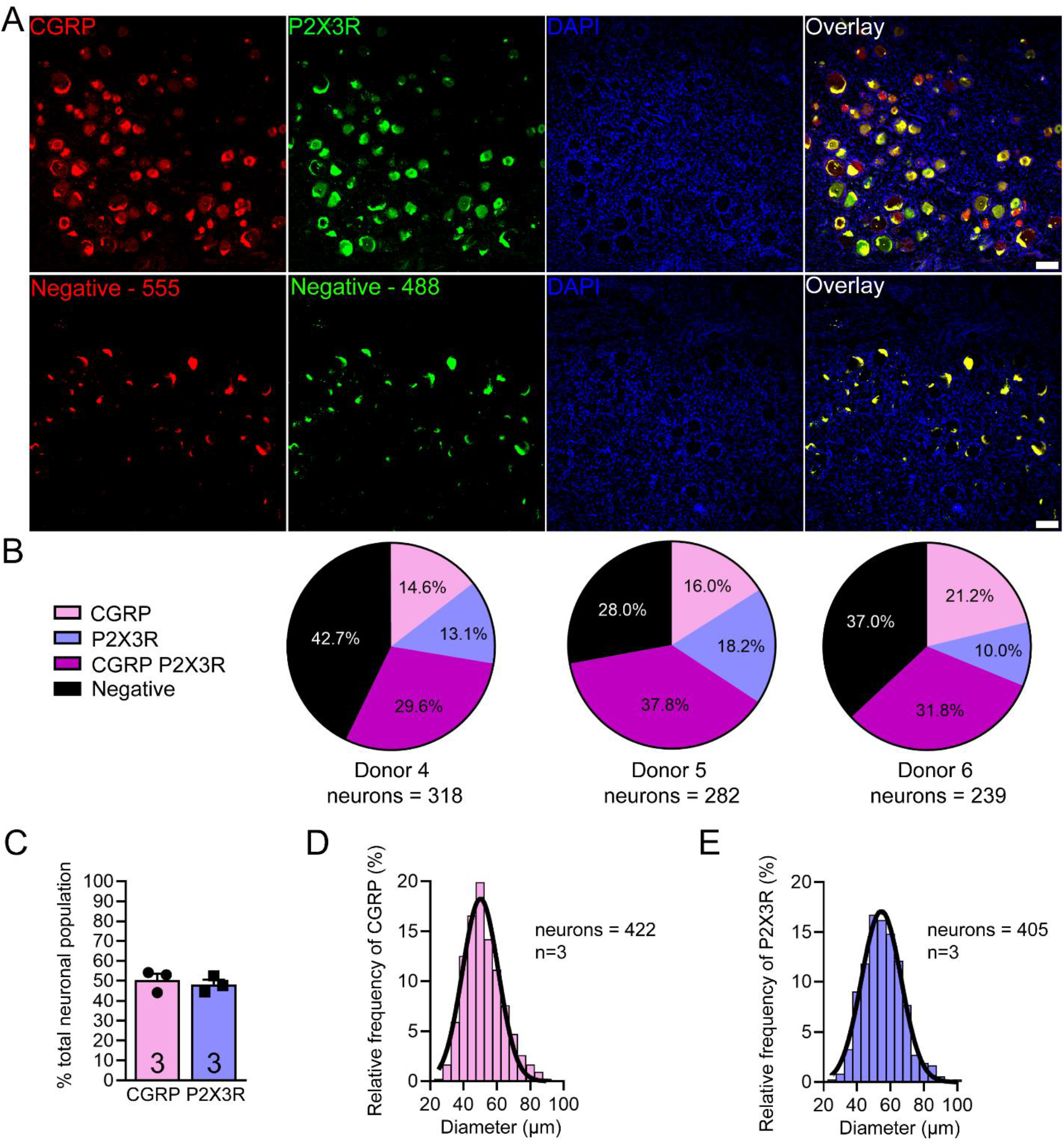
Immunohistochemistry for CGRP and P2X3R in human lumbar dorsal root ganglion. **A)** Representative 10X images showing CGRP (red), P2X3R (green), and DAPI (blue) staining in human dorsal root ganglion (DRG). The negative control was exposed only to secondary antibody and was imaged at the same settings. **B)** Pie-charts showing the distribution of CGRP and P2X3R neuronal subpopulations for each human donor. A large overlapping population of CGRP and P2X3R was observed in all three donors. **C)** CGRP and P2X3R were expressed in 50.4% and 48.3% of all human sensory neurons, respectively. **D)** Histogram with Gaussian distribution displaying the size profile of all CGRP-positive neurons and **E)** P2X3R-positive neurons. DRGs from donors 4, 5 and 6 were used for this experiment. Scale bar = 100μm.

### TrpV1 protein is expressed by most putative nociceptors in human DRG

We next investigated whether TrpV1 protein showed similar expression to its mRNA, which we previously reported was found in ~70% of all sensory neurons in human DRG (Shiers et al., 2020). We used a TrpV1 human-specific antibody that is heavily cited in experiments on human tissues (de Fontgalland et al., 2014; Dinis et al., 2005; Gonzales et al., 2014; Pecze et al., 2008). Again, we observed a gradient of expression for TrpV1 ranging from very low - very high signal (**Fig 2A**). We used the same conservative approach and considered positive neurons as those showing fluorescence intensity signal in the moderate - very high range. With these parameters, TrpV1 protein was found in 57.8% of all sensory neurons (**Fig 2B**) which were also small-to-medium in size (**Fig 2C**). This finding is also consistent with previous findings at the mRNA level in human DRGs from distinct organ donors (Shiers et al., 2020).

**Figure 2.**
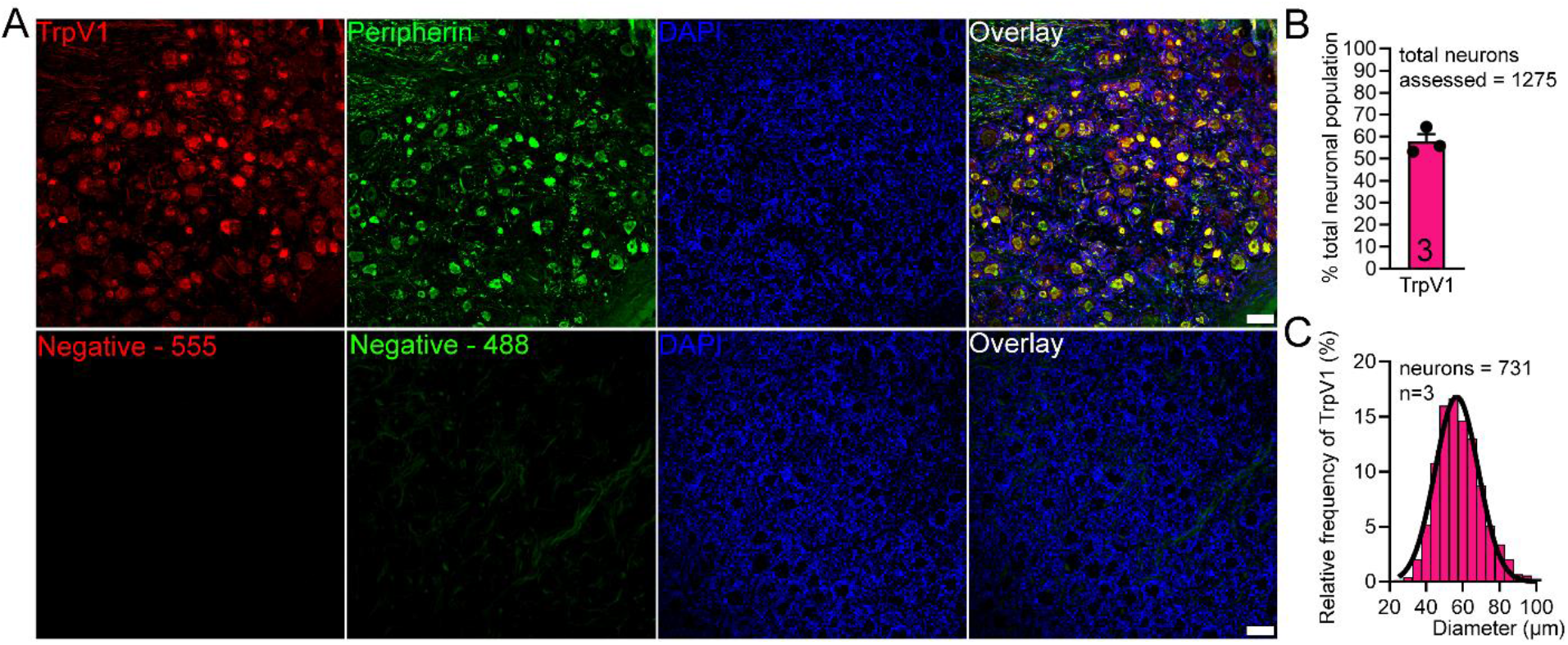
Immunohistochemistry for TrpV1 in human lumbar dorsal root ganglion. **A)** Representative 10X images showing TrpV1 (red), peripherin (a sensory neuron marker; green) and DAPI (blue) in human DRG. The negative control was exposed only to secondary antibody and was imaged at the same settings. **B)** TRPV1 was expressed in 57.8% of all human sensory neurons. The total number of neurons assessed between all three donors is shown on the bar graph. **C)** Histogram with Gaussian distribution displaying the size profile of all TrpV1-positive neurons. DRGs from donors 1, 4 and 5 were used for this experiment. Scale bar = 100μm.

### CGRP, P2X3R, TRPV1, and Nav1.7 protein are present throughout the entire substantia gelatinosa

Next, we assessed the distribution of CGRP and P2X3R protein in human lumbar spinal cord obtained from organ donors. We predicted that these two markers would label similar laminae in the human spinal cord given their high co-expression in human DRG. As noted previously, CGRP and P2X3R label highly distinct populations of neurons in mouse DRG (Shiers et al., 2020) and as such, they label different laminae in mouse spinal cord. In mouse, CGRP innervates lamina I and lamina IIo of the spinal dorsal horn while P2X3R almost exclusively labels lamina IIi (Abrahamsen et al., 2008; Park et al., 2008).

In human, we observed a neuropil staining pattern for both CGRP and P2X3R in the spinal dorsal horn (**Fig 3**). In order to assess the laminar organization of this signal, we conducted H&E staining of adjacent sections of human spinal cord to help visualize the substantia gelatinosa which represents all of lamina II (Rexed, 1952, 1954; Ribeiro-da-Silva, 2015) and aligned it with our IHC images (**Fig 4**). We noted that CGRP and P2X3R labeled the entirety of the substantia gelatinosa but gave little-to-no labeling in lamina I, therefore, we could not define boundaries for that area (**Fig 4**). We also observed that CGRP gave uniform, diffuse staining throughout the entire lamina II region, while P2X3R gave weaker signal in the dorsal portion of lamina II which we defined as lamina IIo, and a stronger signal in the more ventral portion which we labeled as lamina IIi (**Fig 4**). This is similar to observations previously reported in the rat (Ribeiro-da-Silva, 2015). We performed this type of alignment analysis from lumbar spinal cord sections from four organ donors and the data is reported in **Table 3**.

**Figure 3.**
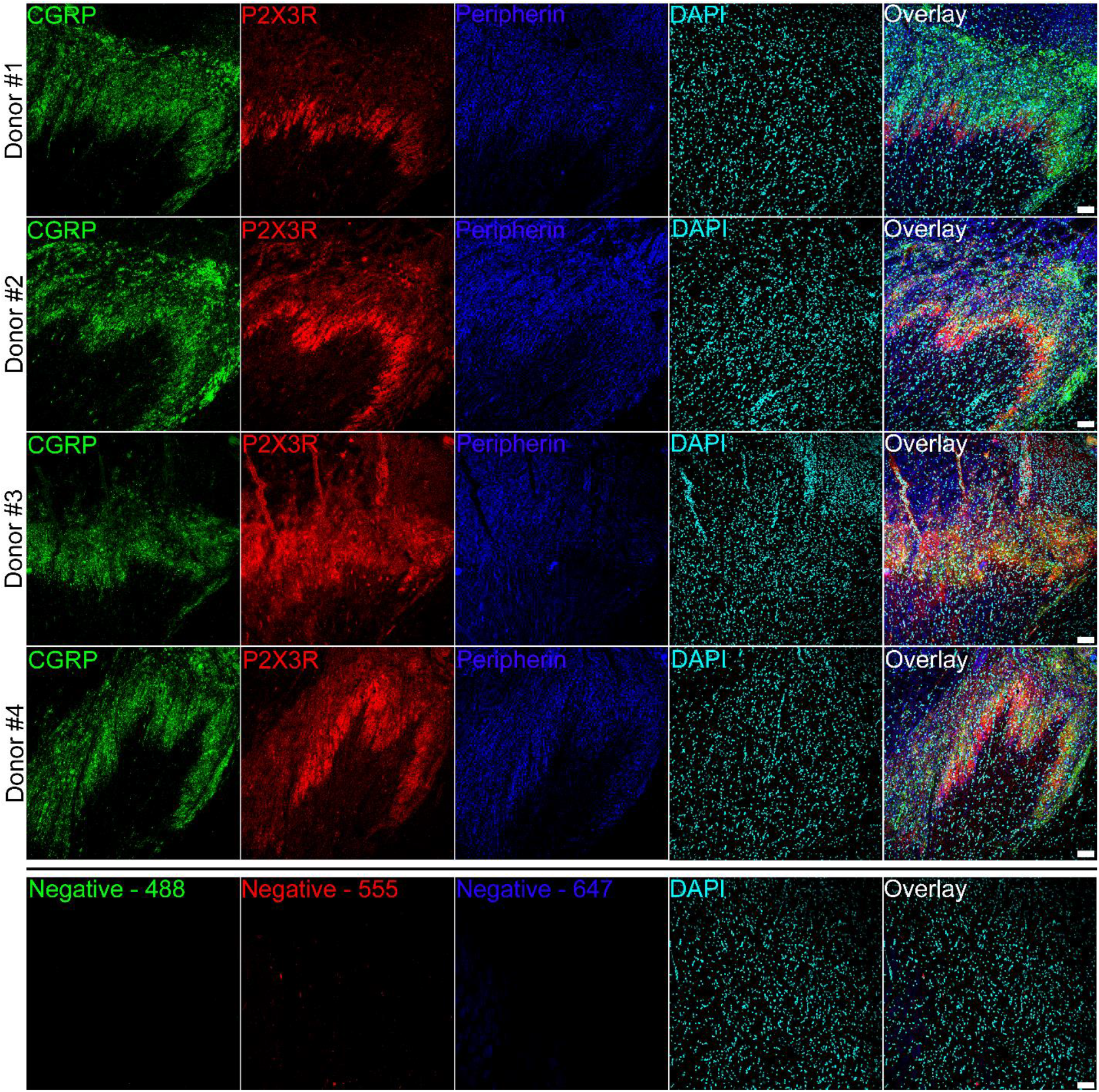
Immunohistochemistry for CGRP and P2X3R in human lumbar spinal cord. Representative 10X images showing CGRP (red), P2X3R (green), Peripherin (blue) and DAPI (cyan) staining in human dorsal root ganglion (DRG). The negative control was exposed only to secondary antibody and was imaged at the same settings. Spinal cords from donors 1, 2, 3 and 4 were used for this experiment. Scale bar = 100μm.

**Figure 4.**
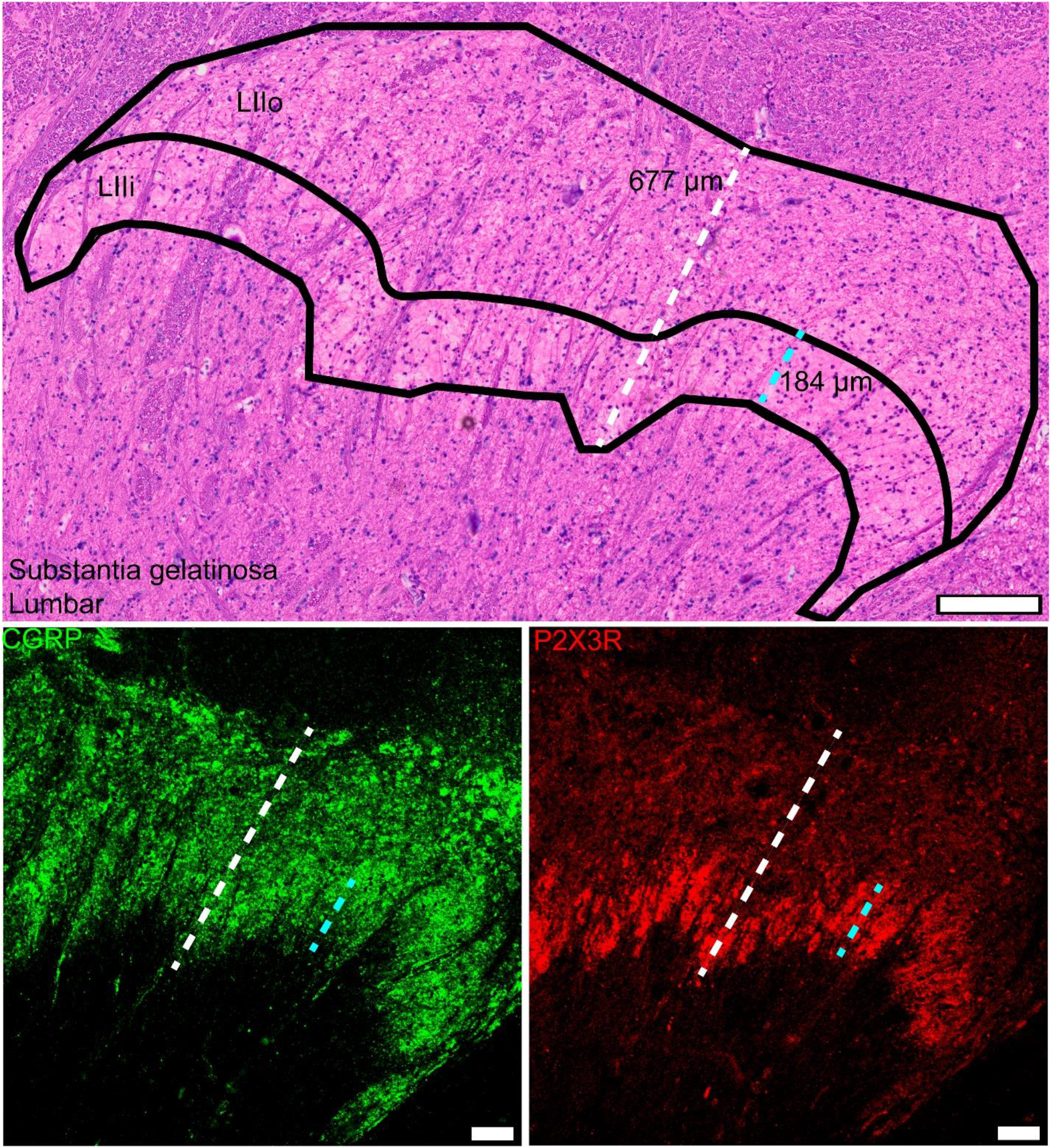
Substantia gelatinosa analysis using Hematoxylin and Eosin (H&E) staining of human lumbar spinal cord. Representative image of an H&E stained section of human lumbar spinal dorsal horn with corresponding CGRP and P2X3R IHC on an adjacent section. Measurements of the entire substantia gelatinosa (transparent, “gel-like” appearance) was made in the H&E stained images from all donors. Carefully constructed guide lines were measured in CellSens in order to measure the same ventral-to-dorsal axis on both sets of the images. Cyan line = dense band of P2X3R label which we denoted as lamina II inner. White line = total length of lamina II. Spinal cords from donors 1, 2, 3 and 4 were used for this experiment. Scale bar = 100μm.

**Table 3.**
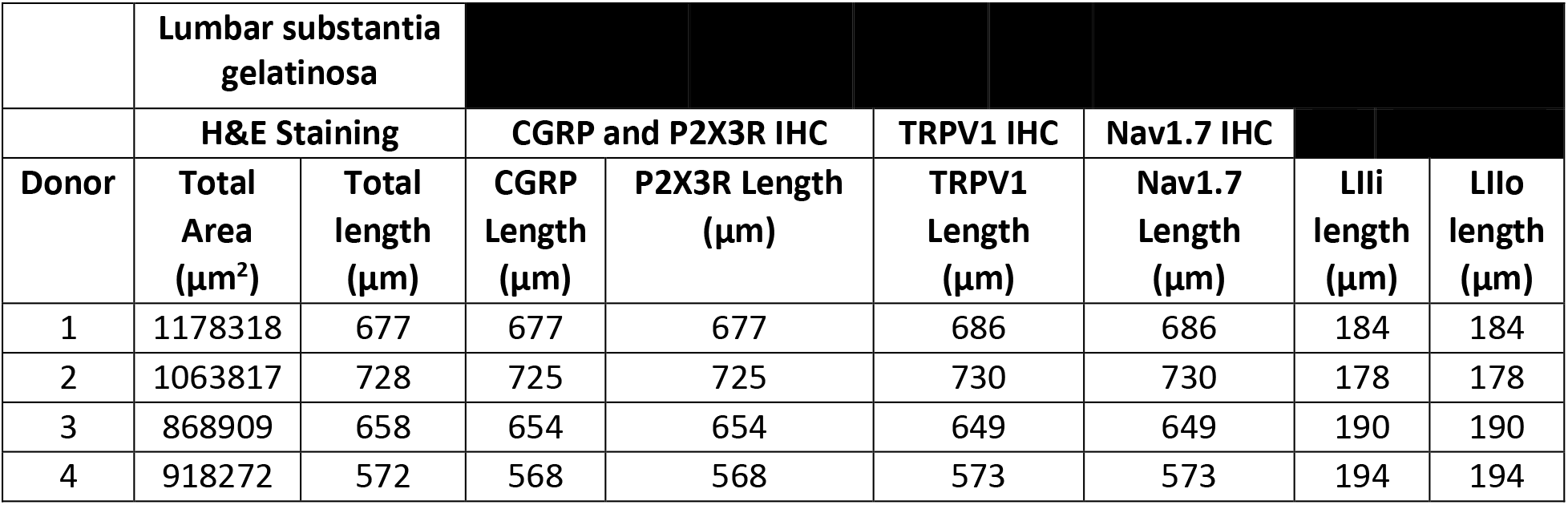
Quantitative measurements of the substantia gelatinosa and CGRP, P2X3R, and TRPV1 IHC label in the human lumbar spinal dorsal horn. The total area of the substantia gelatinosa was measured using H&E staining. The ventral to dorsal length of the substantia gelatinosa was measured in the H&E images and then the same spot was measured on the IHC images (CGRP, P2X3R, TRPV1 and Nav1.7). We generated several length measurements using the lateral aspect of lamina II as our hub point to make sure we were measuring the same spot between both images. The lamina II inner (LIIi) length measurements represent the ventral to dorsal axis length of the dense, strong intensity P2X3R band. Lamina II outer (LIIo) measurements were calculated by subtracting the LIIi length from total length of the substantia gelatinosa.

We executed the same analysis on human spinal cord sections stained for TrpV1 and Nav1.7 (**Fig 5**). Since we saw expression of TrpV1 protein in most putative nociceptors in human DRG, and we have previously reported that Nav1.7 mRNA and protein is expressed in the majority of human sensory neurons (Shiers et al., 2020), we hypothesized that we would observe neuropil signal for both proteins throughout the entire spinal dorsal horn. Like CGRP and P2X3R, we observed dense neuropil label for Nav1.7 throughout the entire substantia gelatinosa (**Table 3**). Interestingly, TrpV1 was robustly expressed in glial cells, most likely astrocytes as has been previously reported in rodent and human (Ho, Lambert, & Calkins, 2014; Ho, Ward, & Calkins, 2012; Kong, Peng, & Peng, 2017; Martins, Tavares, & Morgado, 2014; Miyake, Shirakawa, Nakagawa, & Kaneko, 2015; Roet, Jansen, Hoogland, Temel, & Jahanshahi, 2019; Schilling & Eder, 2009), but it also gave a neuropil pattern throughout all of the substantia gelatinosa. Denser neuropil signal for TrpV1 was observed in laminar IIi.

**Figure 5.**
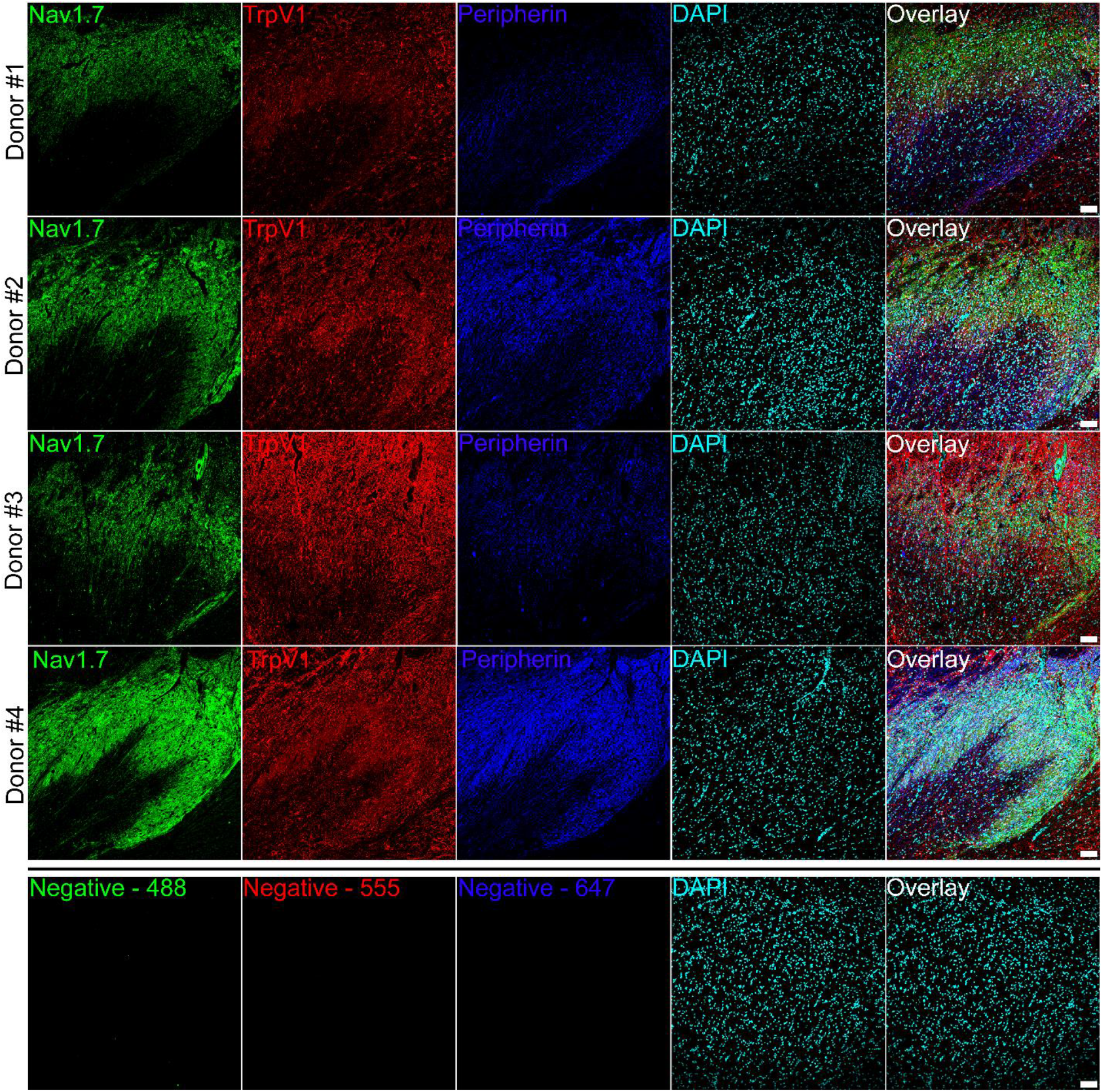
Immunohistochemistry for TrpV1 and Nav1.7 in human lumbar spinal cord. Representative 10X images showing TrpV1 (red), Nav1.7 (green), Peripherin (blue) and DAPI (cyan) staining in human dorsal root ganglion (DRG). The negative control was exposed only to secondary antibody and was imaged at the same settings. Spinal cords from donors 1, 2, 3 and 4 were used for this experiment. TrpV1 and Nav1.7 neuropil staining could be seen throughout the spinal dorsal horn. TrpV1 also robustly labeled glial cells. Scale bar = 100μm.

### Peripherin labels all sensory axons in the human spinal cord

We used peripherin as a co-label for sensory axons in our IHC experiments. We noted that peripherin showed a unique expression pattern in the human spinal cord, labelling axons in the dorsal horn and in the dorsal columns, which contain ascending Aβ fiber axons. To show this, we included mosaic images of peripherin and Nav1.7 staining in human spinal cord to demonstrate the organization of these markers (**Fig 6A**). Peripherin labels the dorsal portion of the entire spinal cord, with particularly strong staining throughout the dorsal horn and dorsal columns (**Fig 6B-C**). Nav1.7 on the other hand is specific to the dorsal horn (**Fig 5 and 6B-C)**. Both can also clearly be seen in incoming axons in the connected dorsal root (**Fig 6D**). High magnification images of these fibers demonstrate the cytoplasmic localization of peripherin within these fibers, and the membranous localization of Nav1.7 (**Fig 6D**). Similarly, high magnification images of the dorsal column show intense peripherin label in Aβ fiber axons with little-to-no Nav1.7 staining (**Fig 6E**).

**Figure 6.**
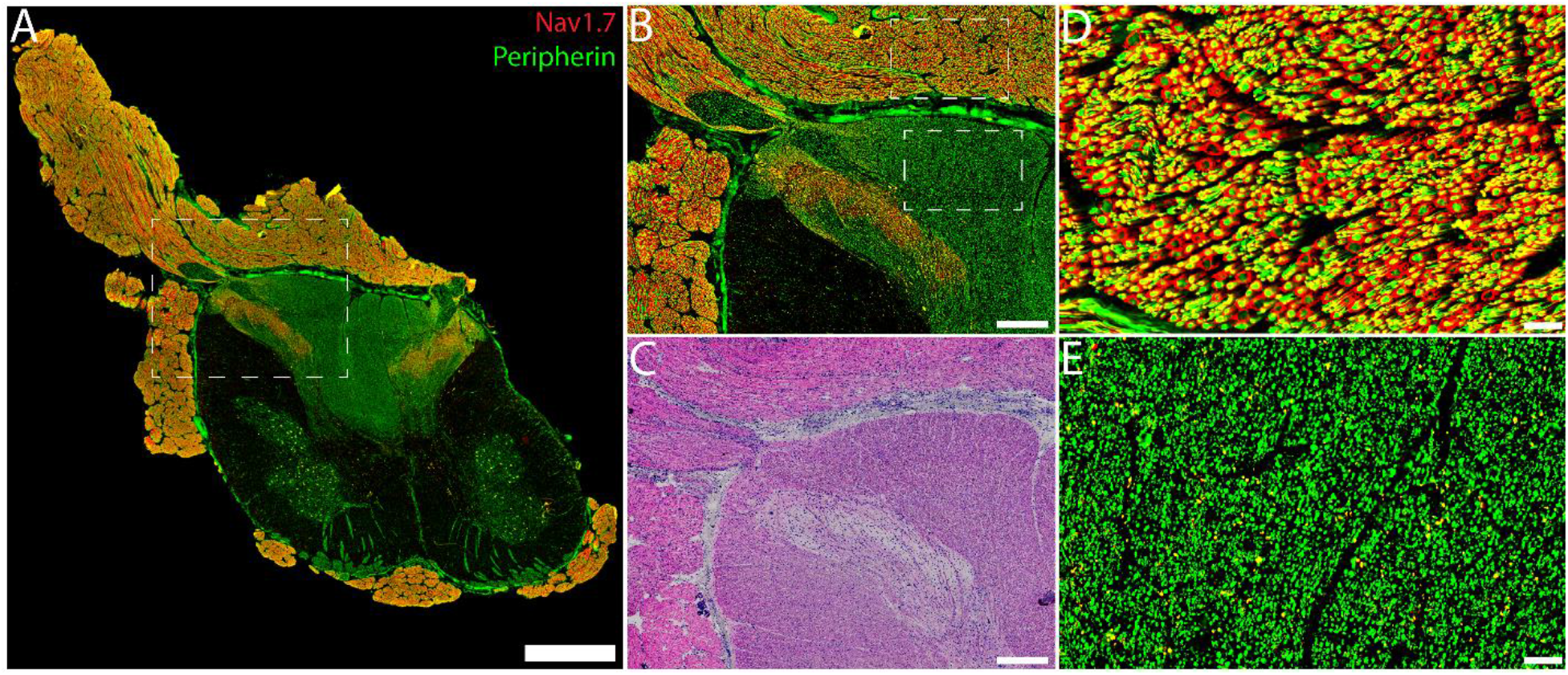
Peripherin labels the dorsal horn and dorsal column of the human spinal cord. **A)** Mosaic image of a human lumbar spinal cord section stained for peripherin (green) and Nav1.7 (red). Scale bar = 2 mm. **B)** Zoom-in of the dashed box in panel A and **C)** the corresponding H&E staining showing the substantia gelatinosa. Scale bars = 500 μm. **D)** Zoom in of the top dashed box in panel B showing Nav1.7 and peripherin signal in sensory neuron fibers in the dorsal rootlet. Nav1.7 appeared to mainly be localized to the axonal membrane while peripherin was found primarily in the axonal cytoplasm. Scale bar = 50 μm. **E)** Zoom in of the bottom dashed box in panel A. Peripherin labeled axons in the dorsal column but little-to-no Nav1.7 signal was detected there. Scale bar = 50 μm.

## Discussion

The use of the terms peptidergic (defined mostly by CGRP expression) and non-peptidergic (defined by IB4 binding and/or P2X3R expression) nociceptors is now almost ubiquitous in the somatosensory and pain fields. This terminology has emerged mostly from work in mice, which are now the most widely used species for basic research in the field (Tuttle, Philip, Chesler, & Mogil, 2018). However, evidence that these populations likely overlap in other species has long existed (Guo et al., 2001; Guo et al., 1999; Perl, 1996; T. J. Price & Flores, 2007; Schmelz et al., 2000). Our present findings, combined with our previous work using *in situ* hybridization techniques (Shiers et al., 2020), as well as many previously published studies (Haberberger, Barry, Dominguez, & Matusica, 2019; T. J. Price & Flores, 2007; Yiangou et al., 2000), clearly demonstrate that peptidergic and non-peptidergic populations are, in the best case, mixed in human DRG. We favor the hypothesis that these segregated peptidergic and non-peptidergic populations do not exist in the human DRG. High throughput data like single cell sequencing will be needed to make direct comparisons between human nociceptors and nociceptors in other species, like mouse.

Peptidergic and non-peptidergic population markers substantially overlap in rat (T. J. Price & Flores, 2007) and human DRG (Shiers et al., 2020) while they are almost entirely segregated in the mouse (Shiers et al., 2020; Usoskin et al., 2015). Other important nociceptor markers like TrpV1 show selective expression at the mRNA and protein levels in a defined subset (~30%) of mouse and rat nociceptive sensory neurons (T. J. Price & Flores, 2007; Shiers et al., 2020); but, in human, the mRNA for *TRPV1* was found in the majority of DRG neurons (~70%) (Shiers et al., 2020) suggesting that it may be expressed by most human nociceptors. However, *TRPV1* mRNA expression may not reflect translated protein. To address this potential difference between mRNA expression and presence of protein, we conducted immunohistochemistry for CGRP, P2X3R, and TrpV1 using human-specific antibodies that have been previously characterized. Like the mRNA pattern, we observed an overlapping population of CGRP and P2X3R-protein expressing neurons in human DRG, and clear indication of TrpV1-expression in most putative human nociceptors. These findings are corroborated by previous studies, such as the previous description of CGRP and P2X3R protein expression in ~60% of human DRG neurons (Nordlind, Eriksson, Seiger, & Bakhiet, 2000; Yiangou et al., 2000). Histology for TrpV1 has been conducted on human DRGs in several studies, but only qualitative analyses were performed (Chang et al., 2018; Facer et al., 2007; Haberberger et al., 2019; Lauria et al., 2006; Li et al., 2015). However, capsaicin responses have been observed in ~60-100% of human sensory neurons cultured from human donors, albeit with a small sample size (Li et al., 2015; Zhang et al., 2019) and microneurography studies have shown that most unmyelinated human nociceptors respond to capsaicin (Schmelz et al., 2000). A larger population of TrpV1 and neurotrophic tyrosine kinase receptor A (TrkA) co-expressing neurons was found in human compared to mouse (Rostock, Schrenk-Siemens, Pohle, & Siemens, 2018).

We also investigated the distribution of CGRP, P2X3R, TrpV1 and Nav1.7 in human spinal cord. To our knowledge this is the first time these markers have been comparatively assessed in the human dorsal horn. The gray matter anatomy of the cat spinal cord was anatomically divided into 10 laminae based on its cytoarchitecture in 1952 by Bror Rexed (Rexed, 1952, 1954). Since then, the Rexed laminae boundaries have been described in other species based on the laminar density of neurons and afferent markers such as CGRP, P2X3R, and TrpV1 (Abrahamsen et al., 2008; Park et al., 2008; Ribeiro-da-Silva, 2015). One of the most easily identifiable features of the Rexed laminae is the existence of the substantia gelatinosa, a transparent “gel-like” region that encompasses all of lamina II (Ribeiro-da-Silva, 2015). We conducted H&E staining and immunolabelling for CGRP, P2X3R, TrpV1 and Nav1.7 demonstrating that these markers are found throughout the entire substantia gelatinosa, suggesting that in the human, neurons expressing these markers project mostly to lamina II. Our findings suggest that lamina I, also known as the marginal zone, may receive fewer direct inputs from nociceptors in the lumbar human spinal cord, that these synaptic contacts are too diffuse to see with the tools we have used, or that these large marginal zone neurons extend dendrites into lamina II to receive synaptic contacts from nociceptors (Cervero & Iggo, 1980). The distribution of peptidergic and non-peptidergic terminals is also different in rat spinal cord with CGRP displaying uniform, diffuse label throughout the majority of lamina I and II and P2X3/IB4 signal throughout all of lamina I and II but with the strongest signal found in lamina IIi (Park et al., 2008; Ribeiro-da-Silva, 2015). These data more closely resemble the staining patterns we observed for CGRP and P2X3R in human spinal cord. CGRP has also been shown to be present throughout the entire dorsal horn of the C1 human spinal cord (Eftekhari & Edvinsson, 2011), but no neuroanatomical co-labels were assessed to differentiate laminar divisions in that study. Similarly, TrpV1 has also been demonstrated throughout the substantia gelatinosa of the human lumbar spinal cord, but no signal for glial cells was noted in this study (Lauria et al., 2006). We observed a similar distribution of TrpV1 protein in human, lumbar spinal cord but we also noted clear signal in glial cells, likely astrocytes (Roet et al., 2019).

While we were not able to investigate synaptic contacts of the afferents described here, our findings support the notion that human lamina II neurons are likely to receive polymodal C and A∂ fiber throughout the outer and inner layers. We believe that this is consistent with previous electrophysiological studies in the cat and monkey. In the cat most lamina II neurons receive inputs from C-fibers, rather than A∂ fibers, but this input is almost always polymodal as the activity of these neurons are modulated by noxious mechanical stimulation and heat (Cervero & Iggo, 1980; Cervero, Iggo, & Molony, 1979). In the rhesus monkey these lamina II neurons, both inner and outer, also receive A∂ inputs but again are polymodal in nature with most neurons responding to both noxious mechanical stimulation and heat (Kumazawa & Perl, 1978; D. D. Price, Hayashi, Dubner, & Ruda, 1979). In macaque monkeys spinothalamic tract neurons in lamina I or II also appear to receive almost entirely polymodal noxious heat and mechanical inputs (Chung, Kenshalo, Gerhart, & Willis, 1979). Our results in the human spinal cord indicate that different subsets of nociceptors project throughout lamina II with no gradient for inner or outer laminae except for afferents marked by P2X3R. While similar studies have not been done in monkey or cat spinal cord, our immunohistochemical findings are consistent with electrophysiological studies described above in those species. A somewhat provocative interpretation of these findings could be that as the spinal cord to brain volume diverges in larger animals (MacLarnon, 1996; Swanson, 1995), processing of nociceptive information occurs more in the brain than in the outer lamina of the spinal cord. This may partially explain why some spinally directed analgesic approaches like NK1 antagonists have failed in the clinic (Hill, 2000).

Why is it important to understand how these population markers, and the populations of neurons themselves, are defined in different species? In our view, the best example of this is that the segregation of these nociceptor populations in the DRG and their synaptic contacts in the dorsal horn has substantiated a labeled line for mechanical and thermal pain which is derived mostly from data in the mouse (J. Braz et al., 2014; J. M. Braz et al., 2005; Cavanaugh et al., 2009; Huang et al., 2019; Mishra & Hoon, 2013). It is now clear from many independent lines of evidence in the mouse that the P2X3R/IB4 population mediates mechanical pain while the CGRP and TRPV1 population mediates heat pain (J. Braz et al., 2014; Cavanaugh et al., 2009; McCoy et al., 2013; Zylka, Rice, & Anderson, 2005). Other labeled lines have been identified in the mouse (Huang et al., 2019; Mishra & Hoon, 2013). While these findings are among the most elegant uses of modern neurobiology behavioral genetics tools, are they relevant to gaining insight into how to treat pain disorders in humans? A major goal of molecular neuroscience on human DRG and spinal cord should be to define the neuronal populations in the ganglia and their synaptic connections in the spinal cord. Our work is a quantitative step in this direction. When this information is thoroughly mapped, we can hopefully then use information gleamed from the mouse to apply what we have learned from interventional studies in that species to better understand how to identify therapeutic targets in the human where such interventional work cannot be done at the discovery stage. This information may also be used to find alternative species for such work where the underlying molecular neuroanatomy may better resemble the human.

## ACKNOWLEDGEMENTS

This work was supported by NIH grants NS065926 and NS111929. The authors are grateful to the organ donors and their families for the gift of life and research provided by their organ donation. We thank Fernando Cervero for helpful discussions on the manuscript.

